# Carbamazepine and GABA have distinct effects on seizure onset dynamics in mouse brain slices

**DOI:** 10.1101/2020.08.11.245951

**Authors:** Dakota N. Crisp, Rachel Parent, Mitsuyoshi Nakatani, Geoffrey G. Murphy, William C. Stacey

## Abstract

Optimizing antiepileptic drug therapy is very challenging due to the absence of a reliable method to assess how brain activity changes between seizures. This work uses the Taxonomy of Seizure Dynamics (Saggio *et al.*, 2020) to investigate how anticonvulsants influence seizure onset dynamotypes. The no Mg^2+^ /high K^+^ mouse brain-slice seizure model (N = 92) was used to generate consistent epileptiform onsets. We compared the onset bifurcations of controls with slices treated with either GABA or carbamazepine. Each anticonvulsant uniquely changed the types of bifurcations in the slices. This experiment provides proof-of-concept evidence that brain states exist on a “map” of seizure dynamics, and that antiepileptic drugs with different mechanisms can change the positioning of the brain states on the map.

**Impact statement:** Antiepileptic drugs modify underlying brain states and influence the pathway into seizure onset in brain slices.

## Introduction

Recent work described a Taxonomy of Seizure Dynamics (TSD), which focuses on how the brain enters and exits seizure states, modeling these transitions using bifurcations (Saggio *et al.*, 2020). The TSD describes four seizure onset bifurcations: Saddle-Node (SN), Saddle-Node on Invariant Circle (SNIC), Supercritical Hopf (SupH), and Subcritical Hopf (SubH), and demonstrated how they can be distinguished visually with high accuracy (Saggio *et al.*, 2020). This methodology determines the dynamotypes of each seizure onset, and identifies the dynamic regime of the brain immediately before each seizure. Thus, this method potentially assesses how different medications influence the interictal brain state.

Utilizing the TSD to classify seizures has provided evidence that brain states change over time and are correlated with seizure onsets. Humans exhibit a range of dynamotypes when monitored over long periods, suggesting that brain states can vary over the period of days or months (Saggio *et al.*, 2020). In a rodent model of epilepsy, the dynamotypes had similar changes during the progression of epileptogenesis in multiple animals, which corresponded to alterations in the response to perturbing stimuli (Crisp *et al.*, 2020b). These results suggest that the brain does not exhibit a single pathway into seizure, and that dynamotype is influenced by a changing interictal brain state. These experimental findings were originally explained using a “map” of brain dynamics (Saggio *et al.*, 2017).

The current work seeks to determine if various anticonvulsants can alter the underlying dynamics. Our hypothesis is that pharmacological perturbations can move the brain state to different areas of the map, depending on their mechanism. For this initial experiment, we utilize a mouse brain-slice seizure model, which allows for robust testing without the confounds of behavioral state. We find that anticonvulsants alter the underlying state and dynamotype, suggesting that brain states exist on a dynamic map and can be moved via pharmaceutical therapy.

## Methods

### Animals

Wildtype 129SvEv mice (1.7-9.8 months old, mean 5.3, std 2.5) were treated in accordance to previous research (Moore et al. 2011). All procedures were approved by the University of Michigan Institutional Animal Care & Use Committee (protocol 00008648). A total of 38 mice (16 male, 22 female) were deeply anesthetized with isoflurane, then decapitated, and brains sliced, resulting in 92 slices total. We performed an *a priori* power analysis (Χ^2^ – contingency table) using a proxy dataset for sample size estimation. The proxy dataset was nearly identical to the data collected here, but used ketosis (prior to animal sacrifice) as the anticonvulsant property.

### Experimental Procedure

Field recordings were conducted in CA3 of the ventral hippocampus from modified horizontal slices (330 um) (Stoop and Pralong, 2000; McKinney *et al.*, 2009). Brain slices were prepared, maintained, and recorded as previously described (McKinney *et al.*, 2009; Moore *et al.*, 2011). Briefly, slices were prepared (Leica VT1000; Wetzlar, Germany) in an oxygenated, ice-cold, sucrose cutting solution (in mM: 206 sucrose, 26 NaHCO_3_, 10 D-glucose, 2.8 KCl, 2 MgSO_4_, 1.25 NaH_2_PO_4_, 1 MgCl_2_, 1 CaCl_2_, and 0.4 ascorbic acid), transferred into oxygenated, room temperature aCSF (artificial cerebrospinal fluid, in mM: 125 NaCl, 25 NaHCO_3_, 25 D-glucose, 2.5 KCl, 1.25 NaH_2_PO_4_, 1 MgCl_2_, 2 CaCl_2_, and 0.4 ascorbic acid), and rested for at least 1 hour. Slices were then transferred to a recording chamber with constant perfusion of warmed (31-32°C) oxygenated aCSF. Baseline activity was recorded for 1 min, immediately followed by a wash of pro-convulsant solution for the next 44 min.

All slices were treated with a pro-convulsant no Mg^2+^ (0 mM) high K^+^ (10.00 mM KCl) aCSF, referred to hereafter as NMHK. Control slices (41/92) received no additional anticonvulsants, while the two experimental groups of slices were perfused with NMHK + 10 μM gamma-amino butyric acid (GABA) (Williamson *et al.*, 2015) (28/92) or NMHK + 50 μM carbamazepine (CBZ) (Dreier *et al.*, 1998) (23/92). The concentrations of anticonvulsants were chosen based on previous research that showed anticonvulsant efficacy, but not total seizure blockage. GABA (a chloride channel agonist) and CBZ (a sodium channel blocker) were chosen to affect distinct anti-epileptic mechanisms. It should be noted that in any given slice, only one experimental condition was chosen (i.e. NMHK / NMHK+GABA / NMHK+CBZ) and only one recording was taken. In other words, multiple conditions were never tested on the same slice, and a single recording from one brain slice is considered one sample.

### Data Analysis

#### Bifurcation Labeling

Methods for a reliable visual classification of seizure dynamics can be found in our prior publication (Saggio *et al.*, 2020). Reviewers marked only seizure onsets, classifying them as either SN/SubH, SNIC, or SupH. SN and SubH categories were combined, as the experimental setup was incapable of recording DC components, disallowing a separation of the two(Saggio *et al.*, 2020). After validation (see Statistics), final “gold standard” labels were created using the majority-vote consensus for each sample. Samples where reviewers could not agree on a classification were not included in the final conclusions (2/92), bringing the total number of samples down to 90.

#### Statistics

We validated that the visual classification could distinguish the different bifurcations utilizing Fleiss Kappa on the raw reviewer markings to assess inter-rater variability as well as a model fitting procedure that compared the majority-vote reviewer labels (excluding samples with no consensus – 2/92) to algorithmic features computed on the raw epileptiform activity, as described in our previous work (Saggio *et al.*, 2020). We tested the differences between the majority-vote bifurcation labels and each experimental group with a Chi Square test.

## Data Availability

All data and associated scripts/text can be found at the University of Michigan’s Deep Blue Library (Crisp *et al.*, 2020a).

## Results

Epileptiform activity was observed in all three experimental conditions. Only slices that developed sustained epileptiform bursting after being exposed to NMHK were included in the analysis. All three onset dynamics (SN/SubH, SNIC, and SupH) were present in each experimental condition (Fig. 1). We validated that there was high agreement (Table 2) between the human reviewers (p < 1e-324, Fleiss Kappa), and their majority-vote labels captured the differences between the algorithmic features (p < 1e-4, permutation test). Having double-validated our reviewer labels, we tested how the dynamotypes changed with anticonvulsants.

**Figure 1.**
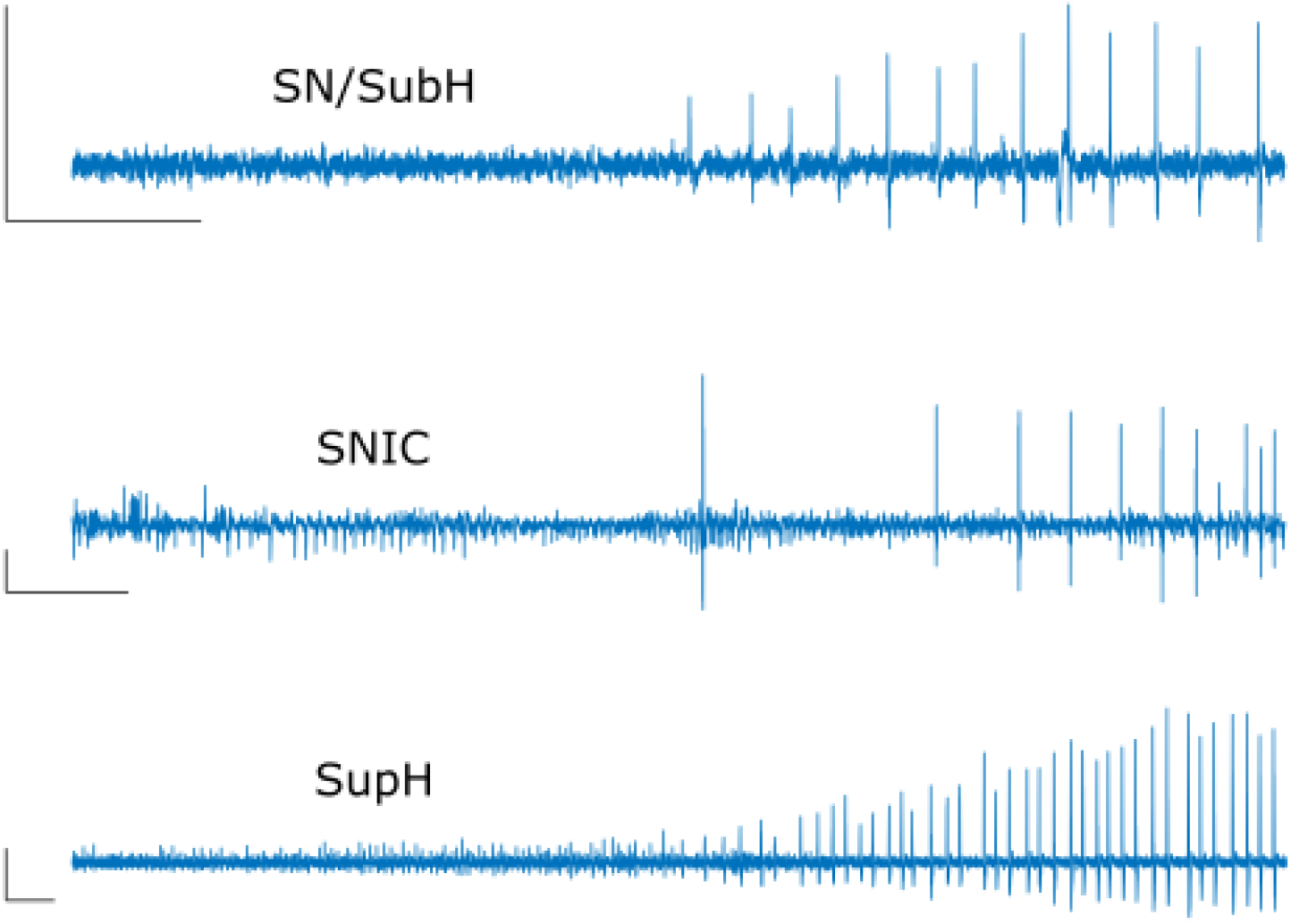
Raw waveform examples of the observed onset bifurcations. All scale bars shown indicate an amplitude scale of 100 mV and a timescale of 10 seconds. Top: a SN/SubH onset is characterized by no specific scaling of amplitude or frequency, here shown as abrupt appearance of periodic spikes. Middle: SNIC onsets appear as full amplitude spikes that increase in frequency. Bottom: SupH onsets have spikes that steadily increase in amplitude from the background noise.

**Table 1.** Rules for visual classification of seizure onset dynamics. Note that SN and SubH cannot be completely differentiated except in the case of a DC shift, which is not always present. A DC shift is indicative of a SN onset. Full description in (Saggio *et al.*, 2020)

Figures
| Visual Condition | Bifurcation Label |
| --- | --- |
| If the amplitude of the spiking activity scales up from 0 | SupH |
| If spikes are increasing in frequency | SNIC |
| If the other two conditions are not met | SN/SubH |

**Table 2.** Reviewer consensus on bifurcation labeling. Reviewers agreed unanimously on the clear majority of seizure onsets (62%). Less than 2.2% of seizures could not be agreed upon by reviewers. Inter-rater variability was computed on all 92 samples. The final analysis comparing onset bifurcation to experimental condition used majority-vote bifurcation labels (N = 90).

|  | Unanimous | 2/3 agree | No consensus |
| --- | --- | --- | --- |
| Onset Bifurcation (N = 92) | 57 | 33 | 2 |

As seen in Fig. 2, in control slices treated only with NMHK, the majority (~63%) of onsets had spikes that started with low frequency and increased over time (SNIC bifurcation). The next most common (~24%) were onsets without any clear trend in ISI or amplitude (SN/SubH), although the spikes were immediately large and distinguishable from baseline. The SupH was the least prominent (~12%). In the slices exposed to GABA (NMHK + GABA), there was a stark shift in dynamics: SN/SubH was most common (~54%), followed by SupH (~35%), and finally SNIC (~12%). In Carbamazepine treated slices (NMHK + CBZ), the most prominent bifurcations were SNIC (~43%) and SupH (~43%). The remaining ~13% were SN/SubH. In summary, onset bifurcations (determined from the majority-vote labels) were found to change significantly between the different experimental conditions (p = 9.1e-5, Chi-squared).

**Figure 2.**
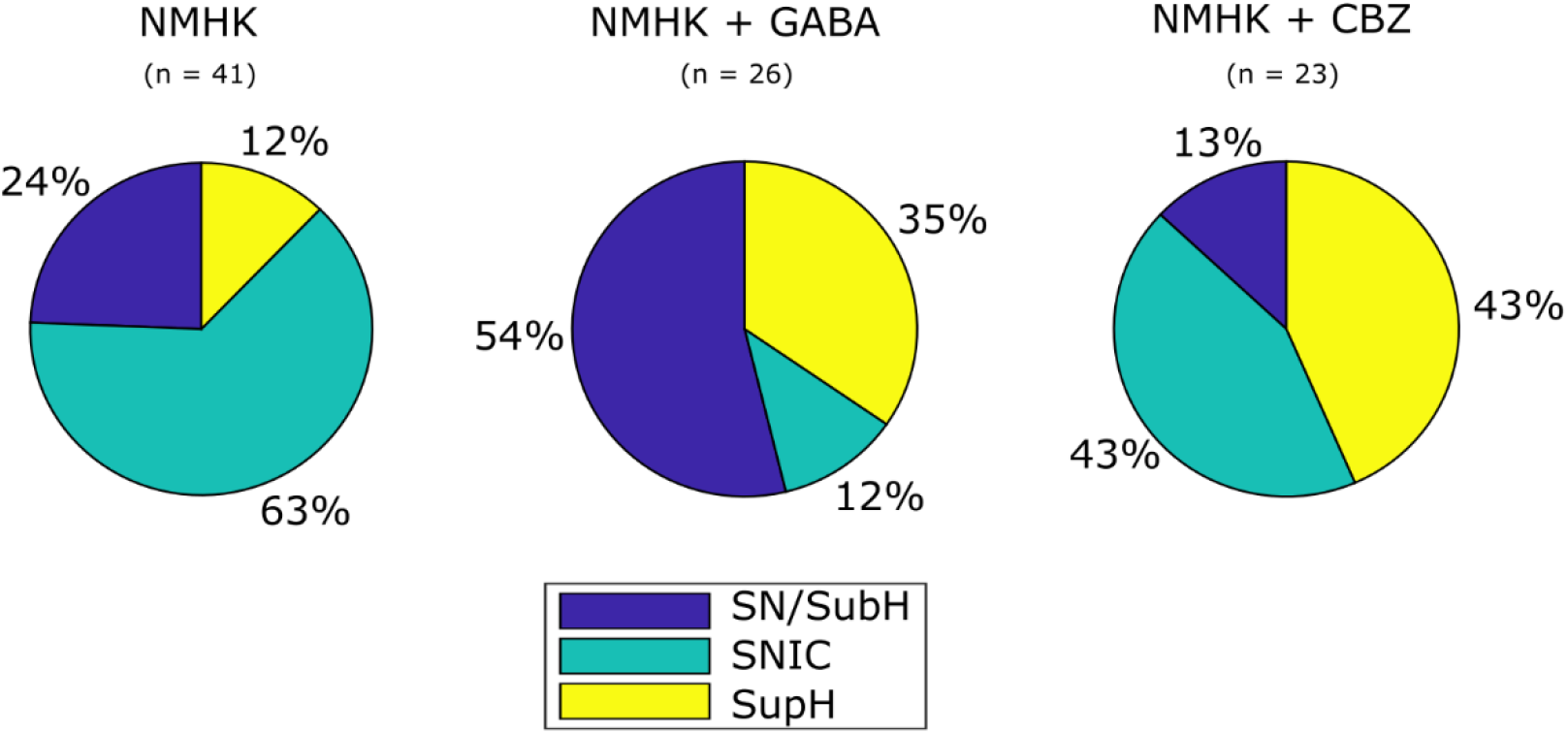
Prevalence of seizure dynamics versus experimental condition. (NMHK) SNIC bifurcations were dominant in slices using only Mg^2+^ /high K^+^. (NMHK + GABA) Introducing GABA decreased the prevalence of SNIC bifurcations, instead increasing the prevalence of both SupH and SN/SubH. (NMHK + CBZ) Introducing CBZ made slices produce equal numbers of SNIC and SupH bifurcations and minimal SN/SubH. Note that in each case, a different bifurcation is significantly less likely.

## Discussion

Using an *ex vivo* model of epileptiform activity, we have discovered that the addition of anticonvulsants can influence how the slice traverses from “normal” to “seizure” state. This experiment was not designed to measure the efficacy of the anticonvulsant—rather, it imposed a highly epileptogenic NMHK solution with subtherapeutic doses of anticonvulsants and assessed dynamotype changes. Both anticonvulsants increased the chance that epileptiform activity started via a SupH bifurcation. And while there were different dynamotypes in each condition, what is most prominent was that one bifurcation group was LESS likely for each pro-convulsant solution. In other words, brain slices without anticonvulsants (NMHK) rarely produced SupH bifurcations; blocking sodium channels (NMHK + CBZ) restricted the production of SN/SubH bifurcations; and chloride channel agonists (NMHK + GABA) restricted SNIC bifurcations. The underlying mechanism(s) remain unclear, however, this study provides evidence that individual anticonvulsants can differentially influence seizure dynamics. These findings could be used to help inform clinical decisions with respect to patient drug selection, and act as further evidence that the epileptic brain exists on a map of seizure dynamics, i.e. that there are certain pathophysiological conditions that tend to place the brain closer or farther away from specific types of seizures, and that this “location on the map” can be manipulated.

The theory of TSD is based purely on the first principles of dynamics, independent of any physical property of the brain. Previous work has shown that vastly different pathophysiology can produce similar dynamotypes (Jirsa *et al.*, 2014), so these results are not meant to show causal relationships. However, this work shows the first evidence that different seizure-promoting conditions can influence the pathway into seizures. These results open the way for future research into how these high-level dynamics can be explained and manipulated by physical properties and interventions.

## Acknowledgements

Funding for this work was provided by the National Institutes of Health (NS094399, AG052934), the University of Michigan EBS Innovation Initiative, and the Horizon 2020 m-Gate project Marie Skłodowska-Curie grant agreement no. 765549. The authors have no conflicts of interest.

